## Supplementary Figures for "Exploration of the genetic landscape of bacterial dsDNA viruses reveals an ANI gap amidst extensive mosaicism"

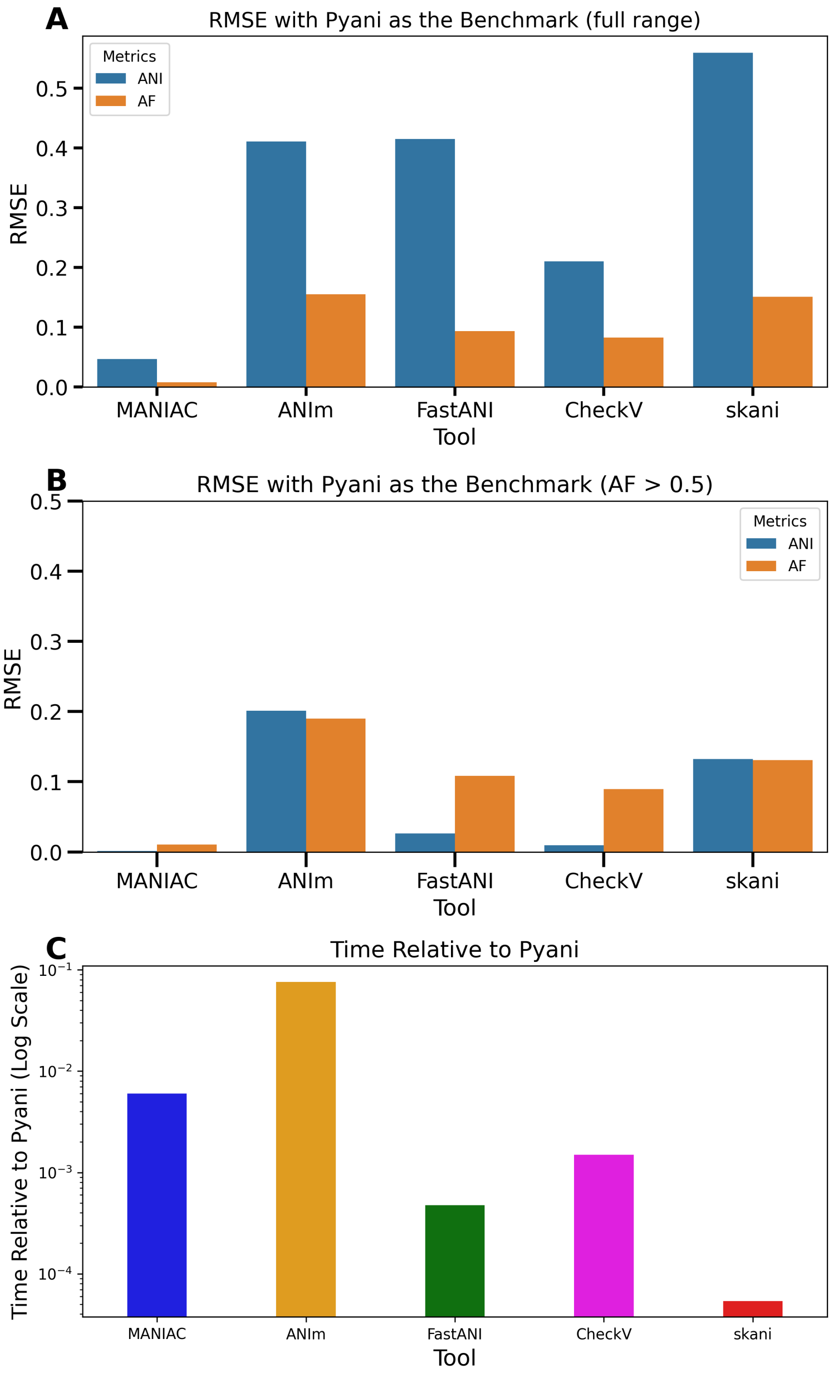


**Figure S1.** (A) Root Mean Squared Error (RMSE) of ANI (blue) and AF (orange) calculation when using ANIb as the benchmark with the same dataset as in Figure 1 (RefSeq 500 dataset) for MANIAC, FastANI, MUMmer, CheckV and skani. (B) Same as in panel A but shown for pairs with AF>0.1 as measured by ANIb. (C) Relative runtime (log10 scale) compared to ANIb when calculating all-by-all distances for different approaches. MANIAC was run with the parameter set **pyani** (see Table 1).

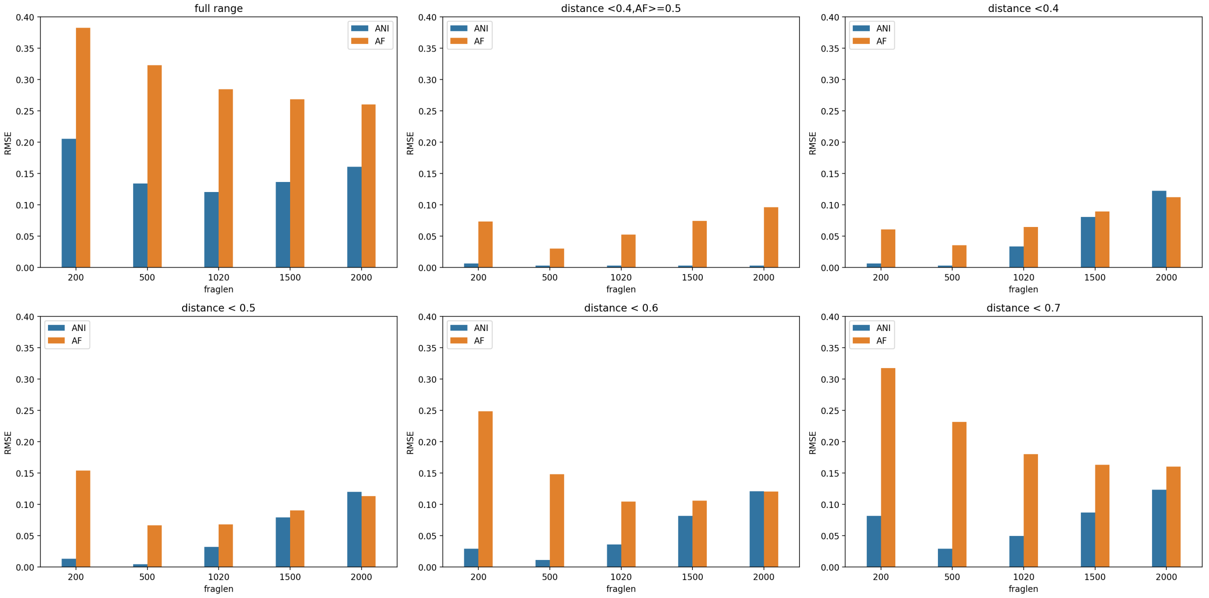


**Figure S2.** Each plot shows root mean square error (RMSE) values for ANI (blue) and AF (orange), obtained by the comparison to the simulated data, calculated for different values of the fragment size: 200, 500, 1020, 1500 and 2000 bp. Panels correspond to different ranges of evolutionary distance $d$ and $c$ over which RMSE is calculated: Full range (Top left), $d < 0.4$ and $c \geqslant 0.5$ (top right), $d < 0.4$ (middle left), $d < 0.5$ (middle right), $d < 0.6$ (bottom left) and $d < 0.7$ (bottom right). All other parameters are provided by the **pyani** paramter set in Table 1.

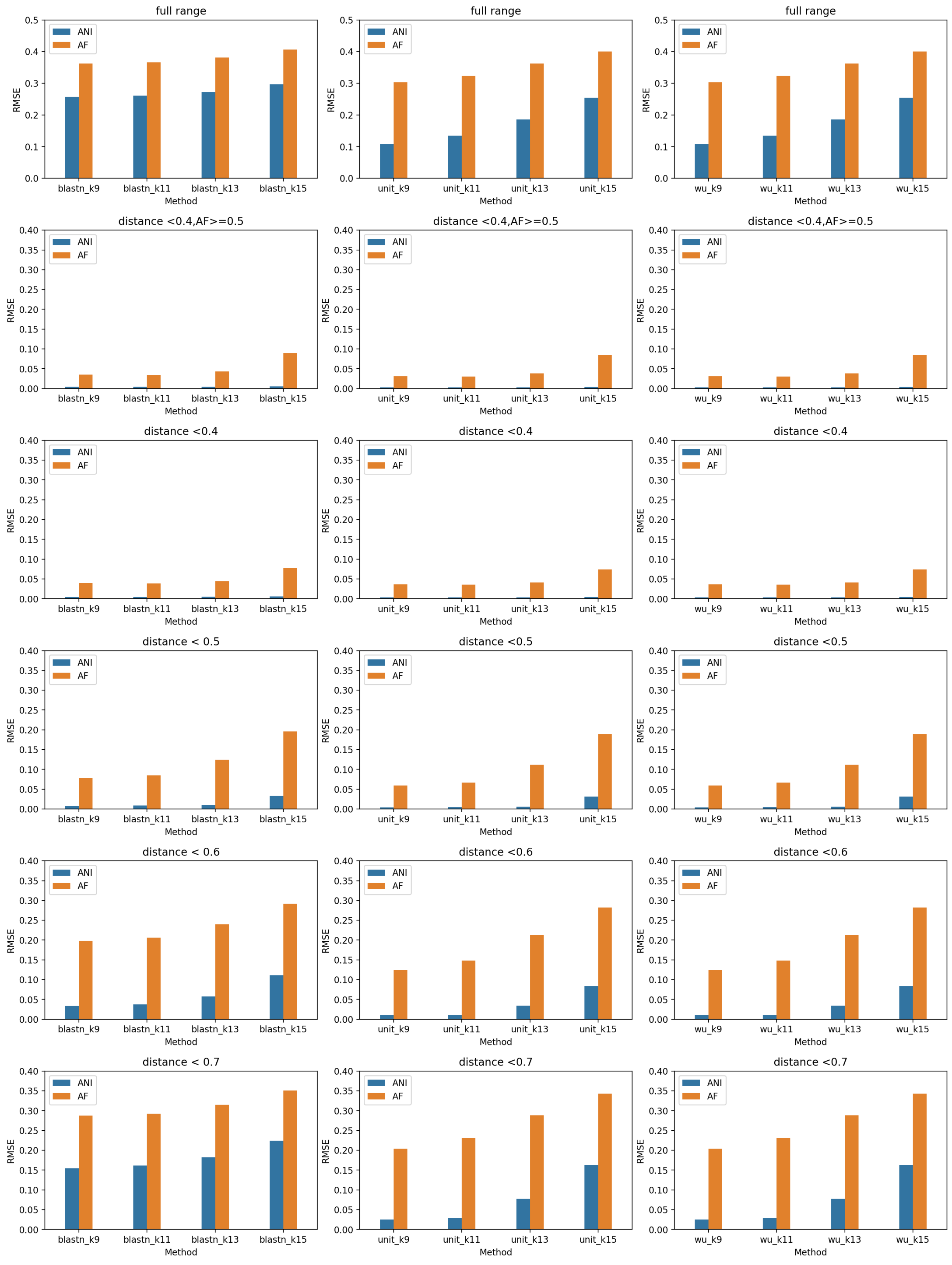


**Figure S3.** Each plot shows root mean square error (RMSE) values for ANI (blue) and AF (orange), obtained by the comparison to the simulated data, calculated for different values of k: 9, 11, 13 and 15. Columns correspond to different scoring schemes: BLASTN (left), UNIT (middle) and WU (right). Rows correspond to different ranges of evolutionary distance $d$ and $c$ over which RMSE is calculated: Full range (first row), $d < 0.4$ and $c \geqslant 0.5$ (second row), $d < 0.4$ (third row), $d < 0.5$ (fourth row), $d < 0.6$ (fifth row) and $d < 0.7$ (sixth row). All other parameters are provided by the **pyani** paramter set in Table 1 with the exception of $F_L$ which was set to 500 as explained in the main text.

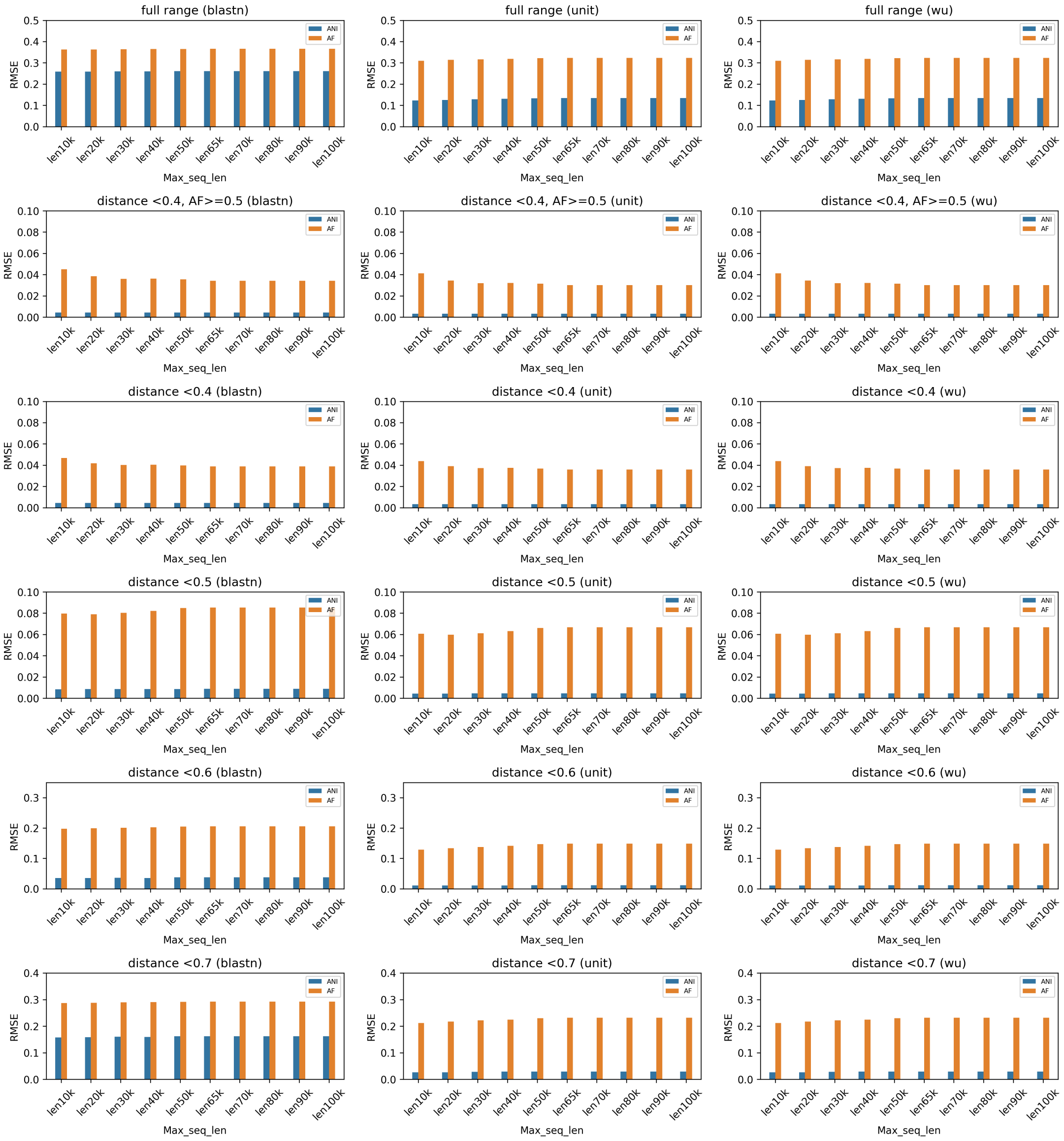


**Figure S4.** Each plot shows root mean square error (RMSE) values for ANI (blue) and AF (orange), obtained by the comparison to the simulated data, calculated for different values of the `max-seq-len` parameter between 10,000 and 100,000, with 65,000 being mmseqs default value. Different plots correspond to different ranges of evolutionary distance $d$ and $c$ over which RMSE is calculated in analogy to Figure S2. All other parameters are provided by the **pyani** paramter set in Table 1 with the exception of $F_L$ which was set to 500 as explained in the main text.

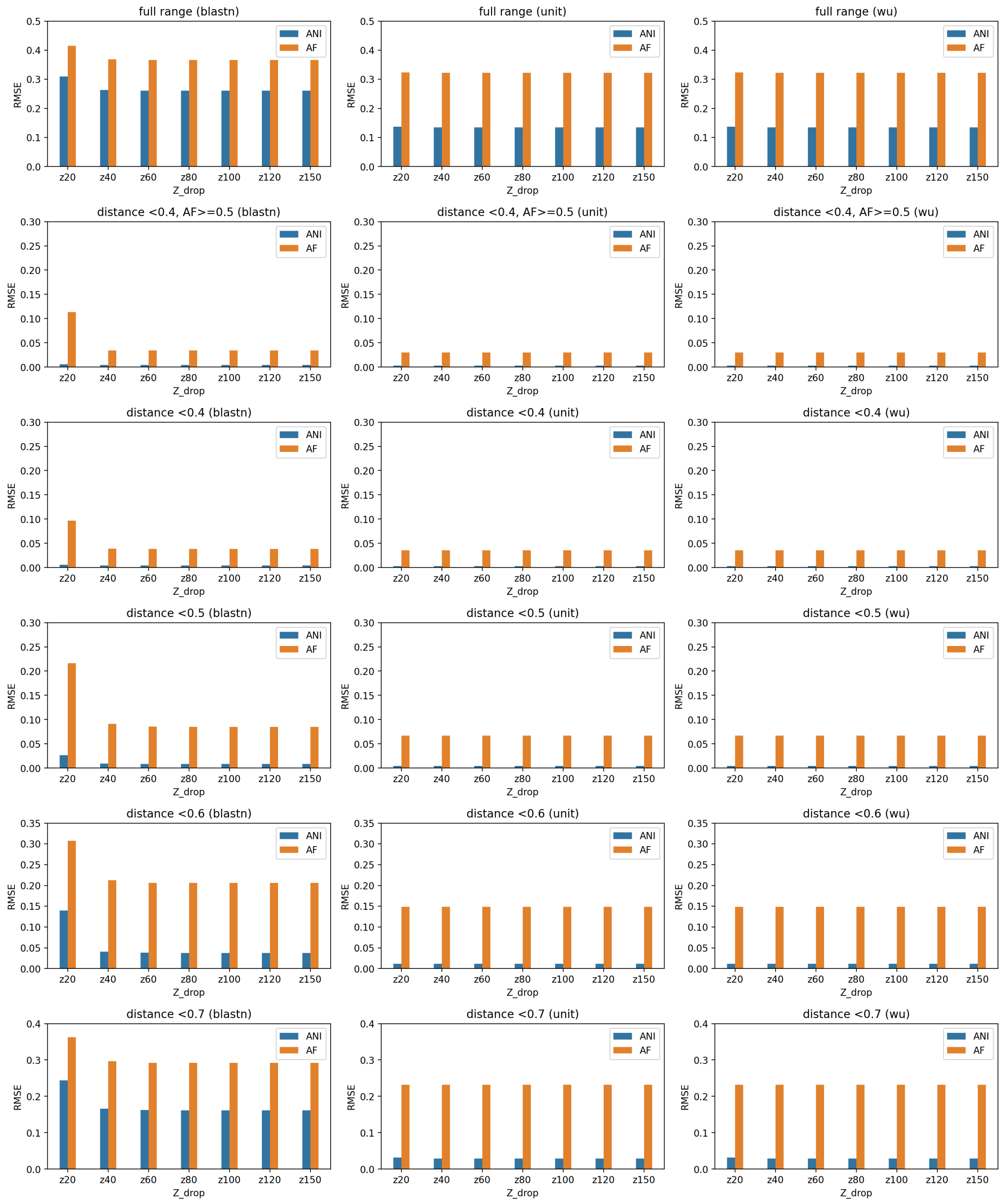


**Figure S5.** Each plot shows root mean square error (RMSE) values for ANI (blue) and AF (orange), obtained by the comparison to the simulated data, calculated for different values of the `zdrop` parameter between 20 and 150, with 40 being mmseqs default value. Different plots correspond to different ranges of evolutionary distance $d$ and $c$ over which RMSE is calculated in analogy to Figure S2. All other parameters are provided by the **pyani** paramter set in Table 1 with the exception of $F_L$ which was set to 500 as explained in the main text.

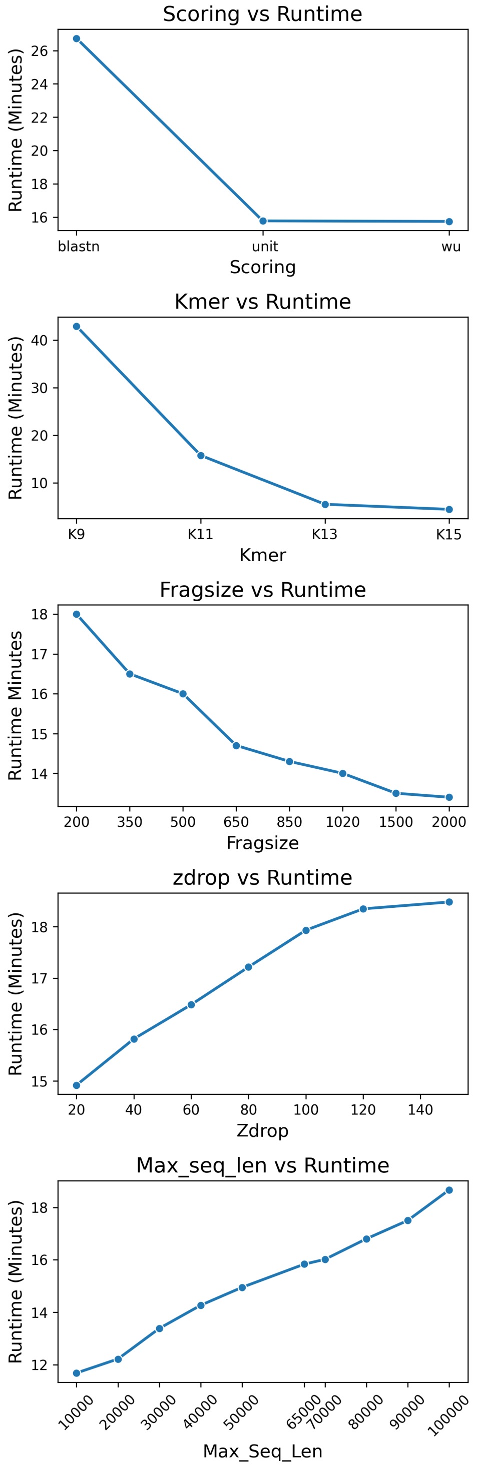


**Figure S6.** Runtimes (minutes) for MANIAC depending on the used parameters: scoring scheme (BLASTN, UNIT, WU), k-mer length (9-15), fragment size $F_L$ (200-2000), `zdrop` (20-150) and `max-seq-len` (10,000 to 100,000). In each plot, the remaining parameters apart from those shown are set to those provided by the **Accurate** paramter set in Table 1.

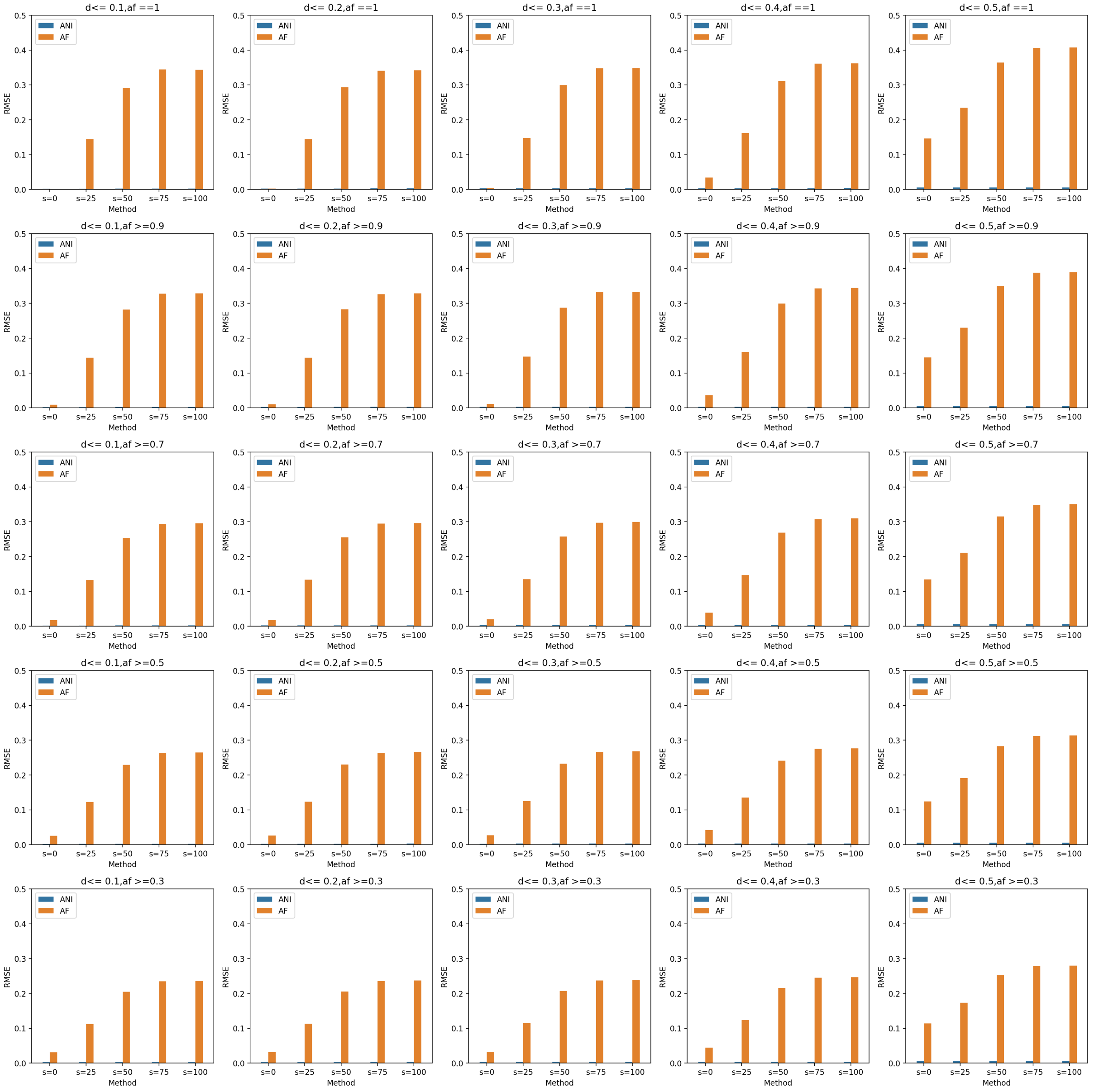


**Figure S7.** Each plot shows root mean square error (RMSE) values for ANI (blue) and AF (orange), obtained by the comparison to the simulated data, calculated for different values of the shuffling parameter, from $s = 0$ to $s = 1$, depending on the range of evolutionary distance $d$ and AF values over which RMSE is calculated. Columns correspond to different ranges of $d$: from $d \leqslant 0.1$ (first column), $d \leqslant 0.2$ (second column), $d \leqslant 0.3$ (third column), $d \leqslant 0.4$ (fourth column), $d \leqslant 0.5$ (fifth column); rows correspond to different ranges of AF: from AF $= 1$ (first row), AF $\geqslant 0.9$ (second row), AF $\geqslant 0.7$ (third row), AF $\geqslant 0.5$ (fourth row), AF $\geqslant 0.3$ (fifth row). We assumed the **Accurate** paramter set in Table 1.

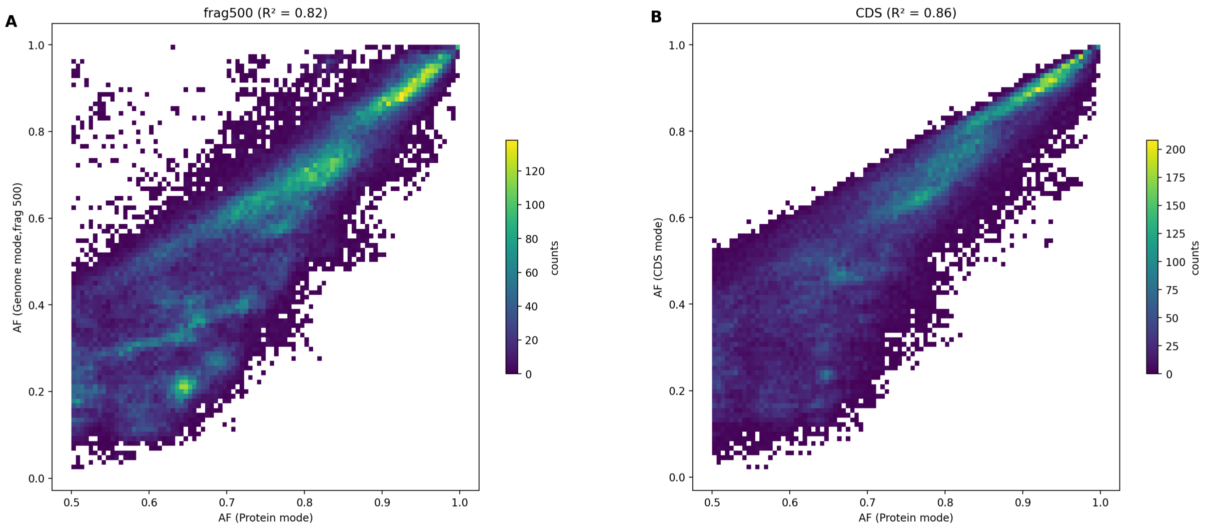


**Figure S8.** Density plot of AF estimated for NCBI RefSeq by MANIAC in Protein Mode (X-axis) vs. AF estimated by MANIAC in Fragment Mode (Y-axis; panelA) or CDS Mode (Y-axis; panelB). R-squared coefficients for linear regression between the two AF values are reported in the corresponding panel. Only pairs with AF $\geqslant 0.5$ (Protein Mode), ANI $\geqslant 0.7$ (Genome Mode, fragment size of 1020bp) and minimal alignment length of 5000 amino-acids were considered.

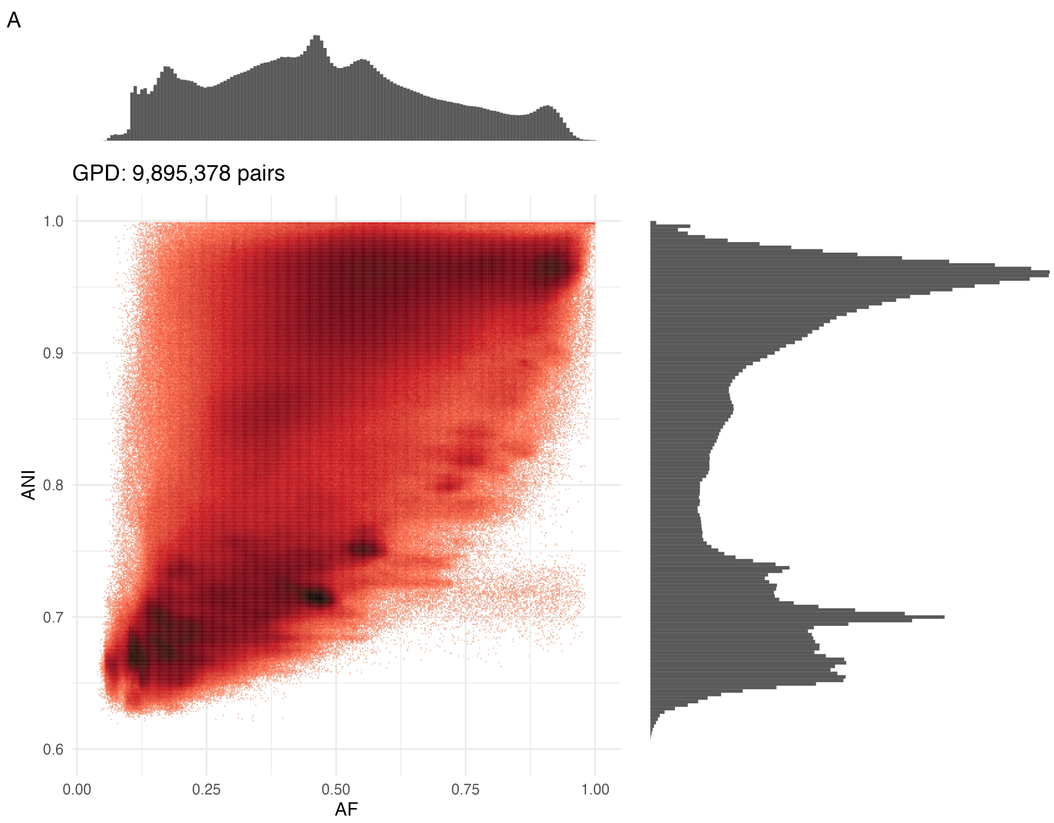


**Figure S9.** Density plot showing the relationship between ANI (Y-axis) and AF (X-axis) for 9,895,378 pairs from the GPD dataset with a minimum alignment length of 10,000 bp. Results are obtained with MANIAC Fragment Mode (both ANI and AF) using the Accurate parameter setting.

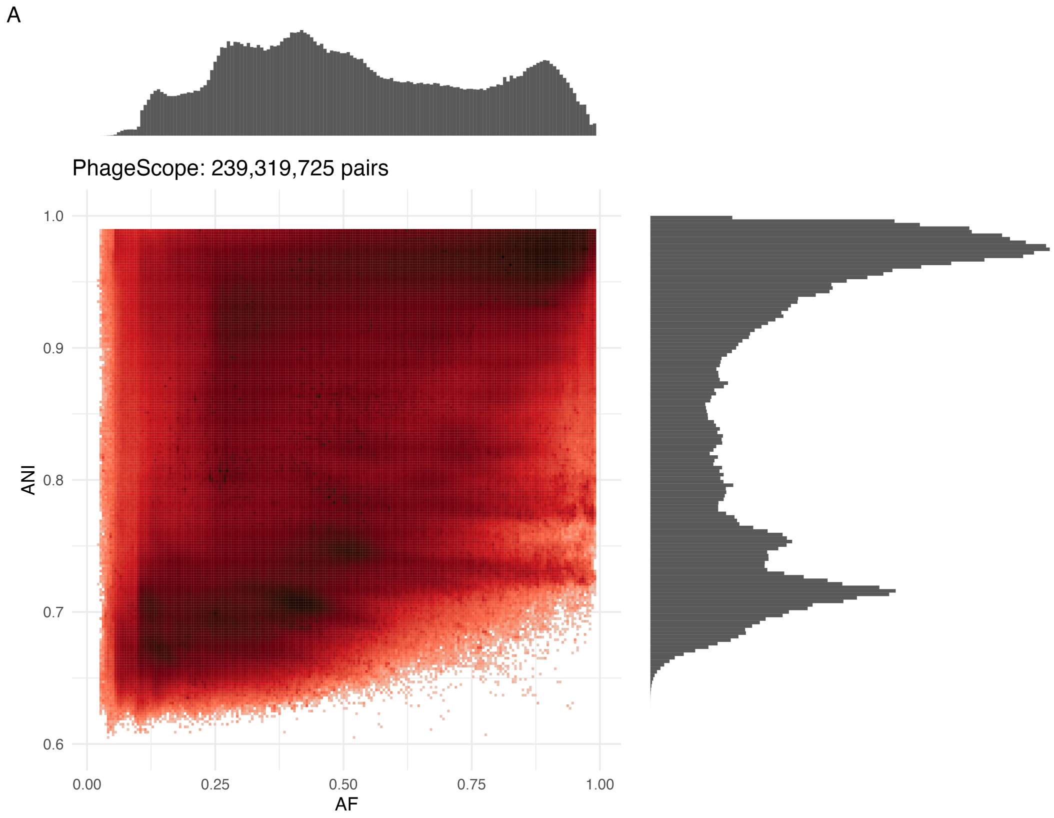


**Figure S10.** Density plot showing the relationship between ANI (Y-axis) and AF (X-axis) for 239,319,725 pairs from the PhageScope dataset with a minimum alignment length of 10,000 bp. Results are obtained with MANIAC Fragment Mode (both ANI and AF) using the Fast parameter setting.

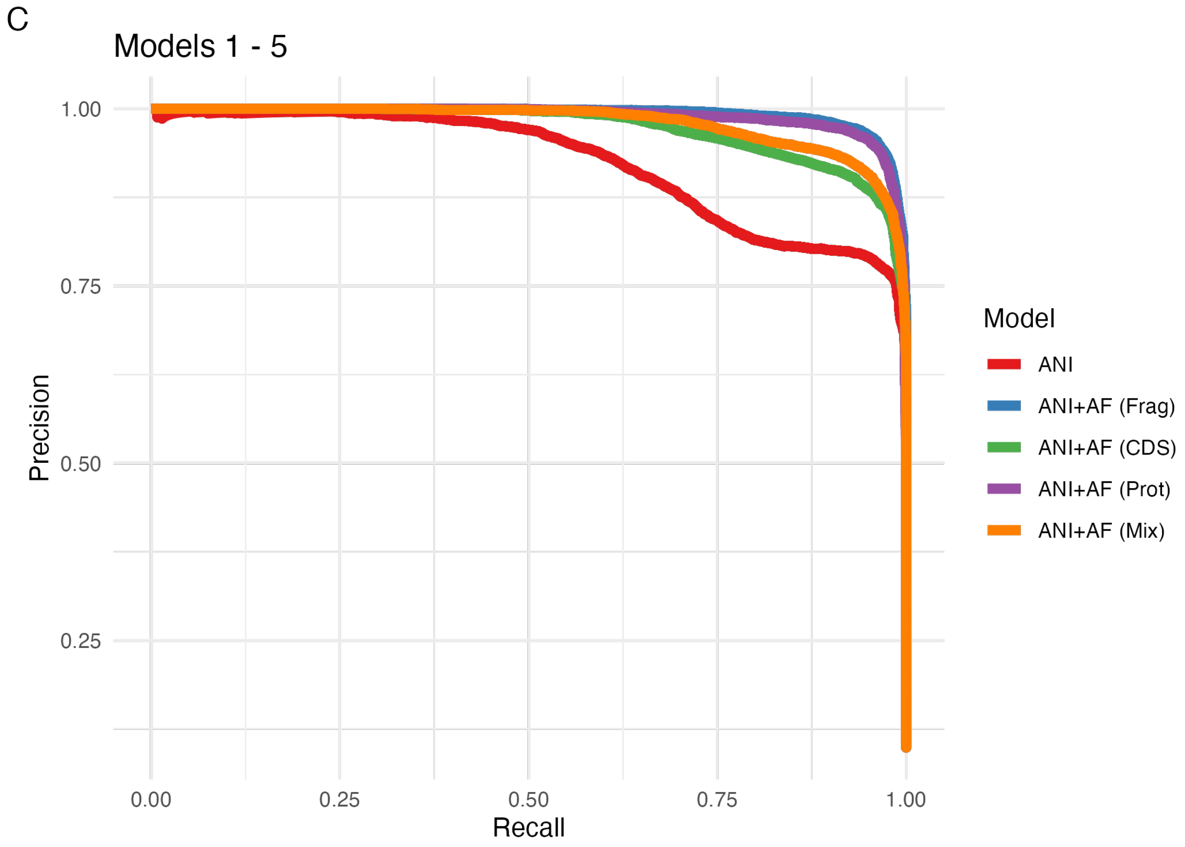


**Figure S11.** PR-AUC curves for the five main models from Figure 6C, obtained when the training/validation and test datasets were split based on the family (instead of genus) assignment, resulting in the 1,047 genera in the training/validation dataset and 999 genera in the test dataset (rest was missing taxonomic assignment at the genus level). The approach was otherwise identical to when datasets were split based on the genus assignment.

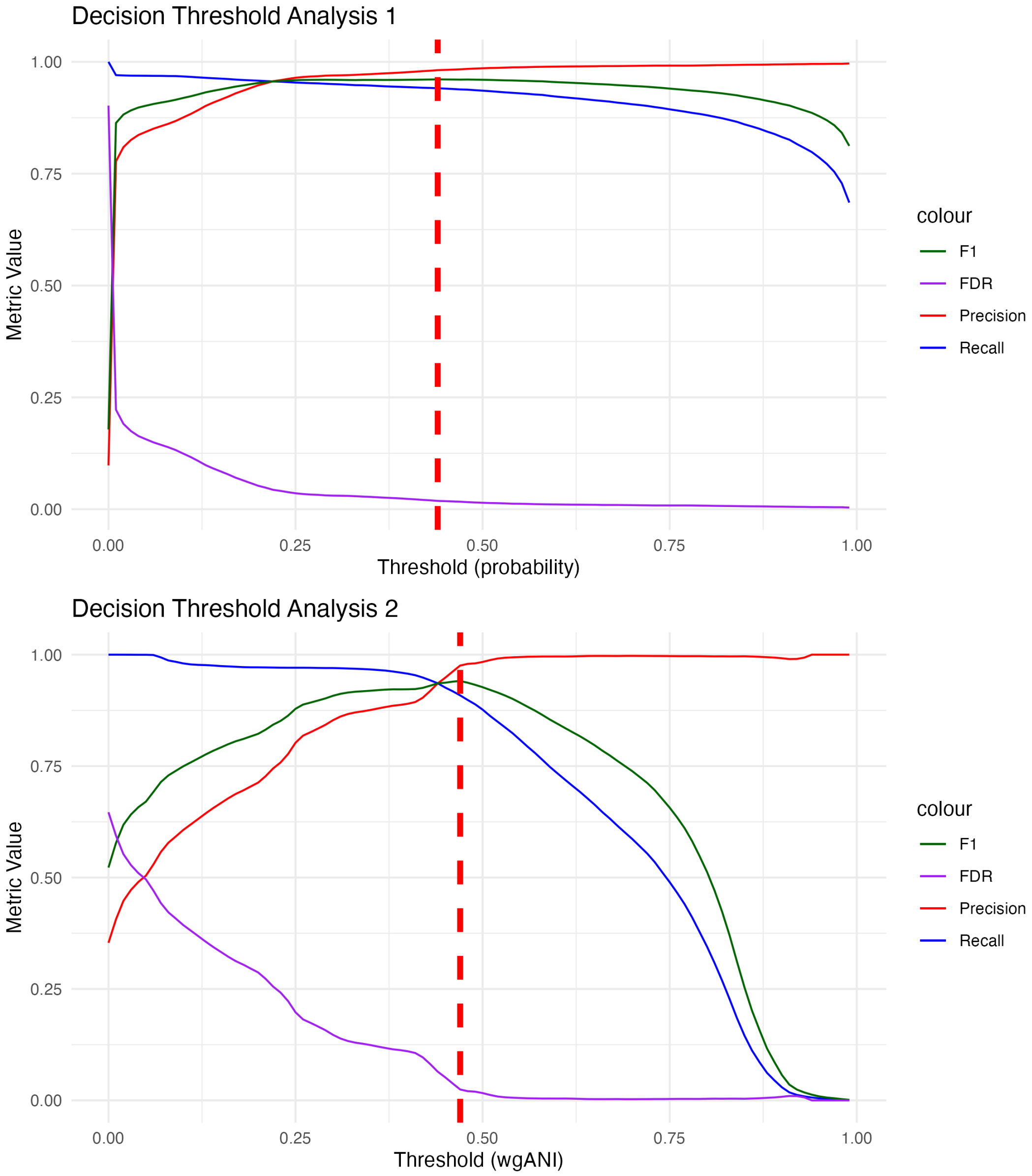


**Figure S12.** Decision threshold analysis. TOP: Precision (red), recall (blue), FDR (purple) values and F1-score (green) as a function of the probability threshold for model 2 (ANI+AF in Fragment Mode) on the test dataset. Red dashed like shows the value ($p = 0.44$) which optimises the F1 score. BOTTOM: Same as TOP but as a function of wgANI threshold, with the value of wgANI$= 0.47$ found to maximise the F1 score. (FDR = false-detection rate).

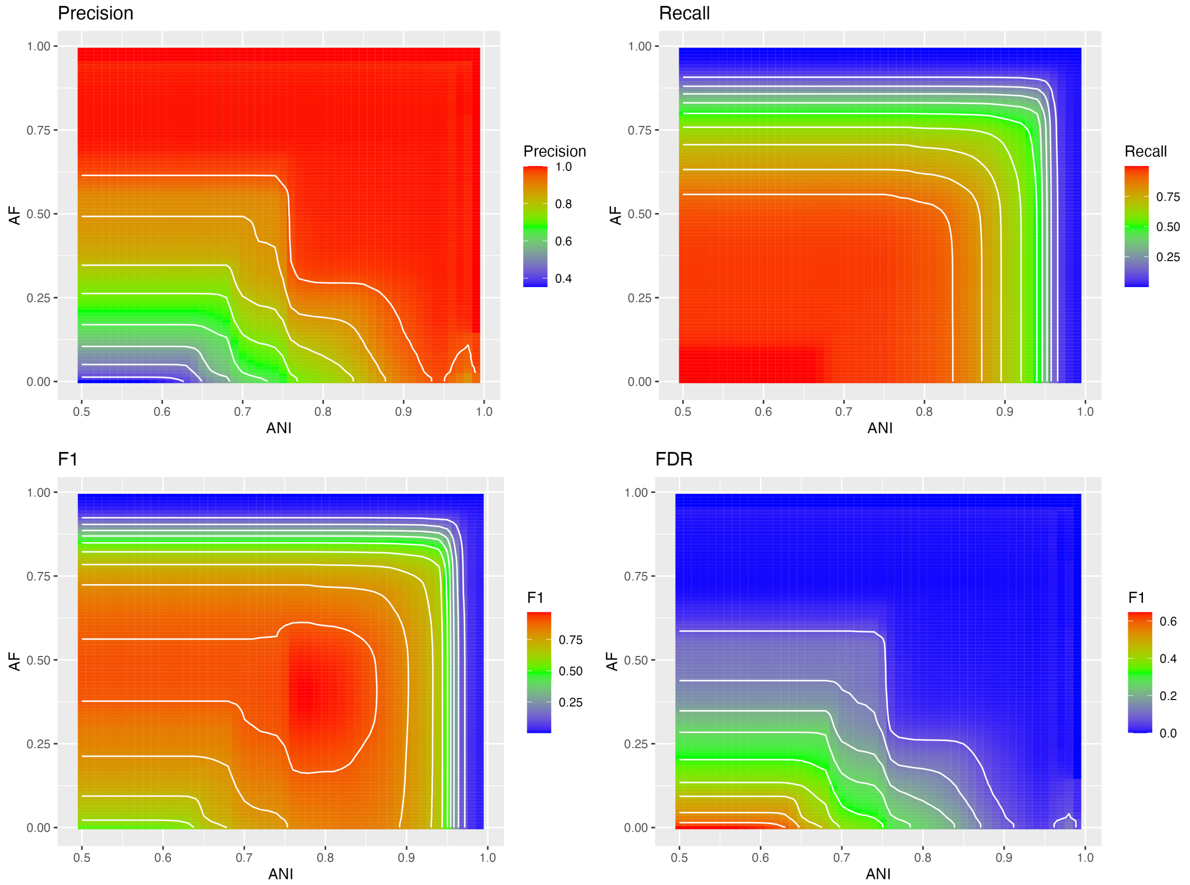


**Figure S13.** Contour plots of precision (A), recall (B), F1 score (C) and FDR (D) depending on the ANI and AF thresholds (calculated in Fragment Mode) to predict same-genus pair on the ICTV dataset. The values were obtained on the test dataset.

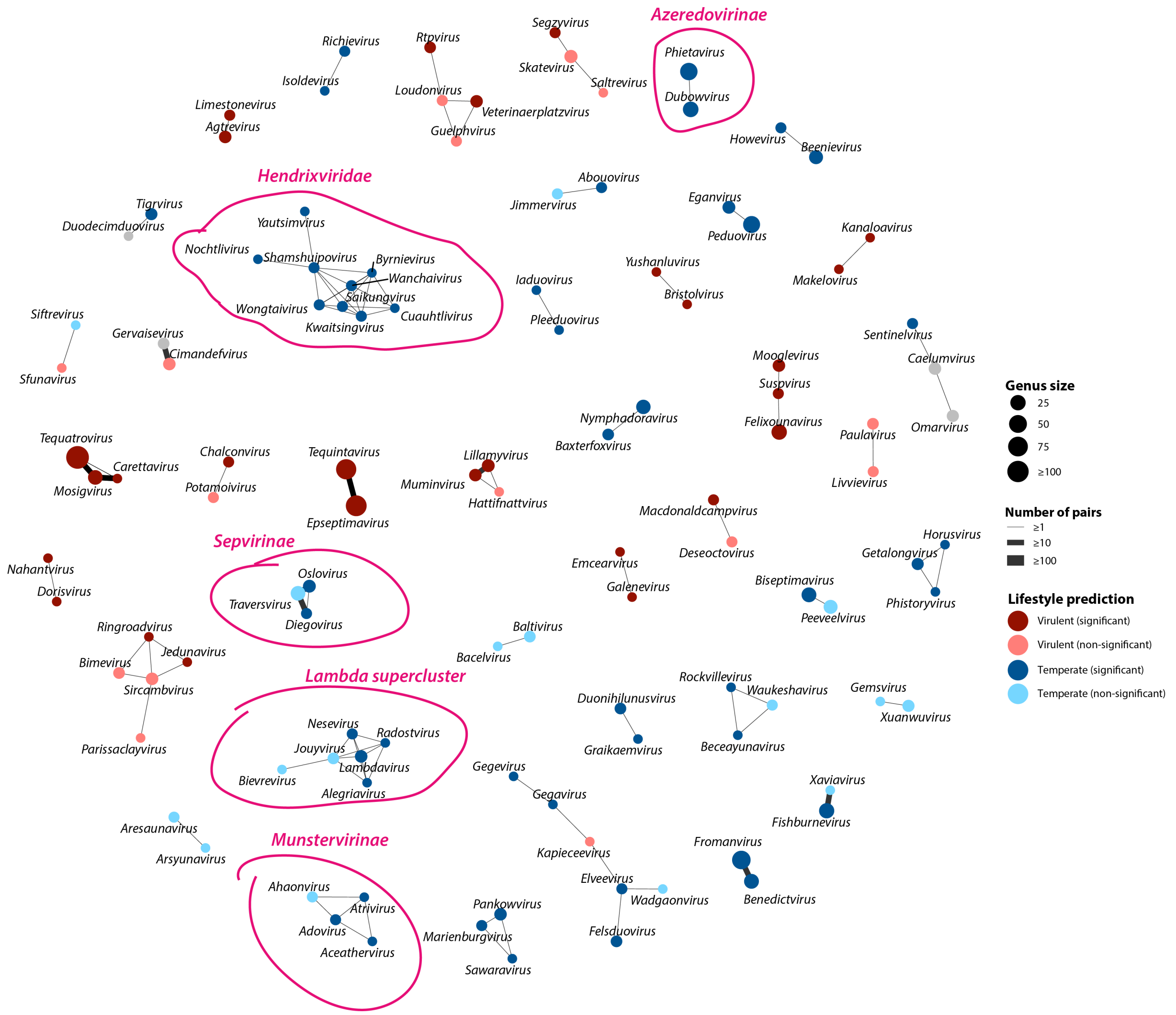


**Figure S14.** Networks shows pairs of 102 viral genera which were predicted to be of same genus with Model 2 (ANI + AF in Genome Mode) with probability $p \geqslant 0.95$ using the entire ICTV dataset (898,323 pairs). Edge weights scale with the number of times the pair was found and predicted in the test dataset. Node size is scaled by the genus occurrence in ICTV39. Colour of the node reflects the predicted lifestyle assignment: red denotes virulent, blue denotes temperate and grey denotes lack of prediction; dark blue/red refers to predictions which are statistically significant ($p \geqslant 0.95$) and light blue/red refers to non-significant predictions ($0.95 > p \geqslant 0.75$).
